# Frequency-specific brain dynamics related to prediction during language comprehension

**DOI:** 10.1101/399733

**Authors:** Kristijan Armeni, Roel M. Willems, Antal van den Bosch, Jan-Mathijs Schoffelen

## Abstract

The brain’s remarkable capacity to process spoken language virtually in real time requires fast and efficient information processing machinery. In this study, we investigated how frequency-specific brain dynamics relate to models of probabilistic language prediction during auditory narrative comprehension. We recorded MEG activity while participants were listening to auditory stories in Dutch. Using trigram statistical language models, we estimated for every word in a story its conditional probability of occurrence. On the basis of word probabilities, we computed how unexpected the current word is given its context (word perplexity) and how (un)predictable the current linguistic context is (word entropy). We then evaluated whether source-reconstructed MEG oscillations at different frequency bands are modulated as a function of these language processing metrics. We show that theta-band source dynamics are increased in high relative to low entropy states, likely reflecting lexical computations. Beta-band dynamics are increased in situations of low word entropy and perplexity possibly reflecting maintenance of ongoing cognitive context. These findings lend support to the idea that the brain engages in the active generation and evaluation of predicted language based on the statistical properties of the input signal.

## 1 Introduction

The brain’s remarkable capacity to process spoken language virtually in real time requires fast and efficient information processing machinery. The efficiency of the brain network for language might in fact rely partially on its ability to dynamically generate predictions from, and apply them to the incoming linguistic signal. In recent years, numerous theoretical accounts of brain function have emphasized the capacity of the brain to use its ongoing contextual embedding for anticipating the future states of environment and the outcomes of its own actions (Bar, 2009; Clark, 2013; Friston, 2005).

The notion of prediction in the brain and probabilistic processing under uncertainty has received increased attention in domains such as memory, perception, and decision-making (Bach & Dolan, 2012; den Ouden, Kok, & de Lange, 2012; Hasson, 2017; Pouget, Beck, Ma, & Latham, 2013). Probabilistic predictive computations in language comprehension, however, have proven to be notoriously elusive to operationalize. As a consequence, the term “prediction” has received differing interpretations in the broader community (Huettig & Mani, 2016; Kuperberg & Jaeger, 2016). In the current study, we turn to computational linguistics, specifically probabilistic language models and information theory, which provide a principled framework for operationalizing predictions in language comprehension as expectation-based processing.

A language model is an estimate of a probability distribution over words in a sentence given a fixed number of words seen so far. Specifically, trigram models used presently estimate the probability of the current word’s occurrence based on the two words just seen. From estimates of probabilities, it is straightforward to compute information-theoretic measures—word perplexity (surprisal) and word entropy—that quantify the amount of information-processing work (in the sense of Shannon information) performed by the cognitive system in transitioning from one word to the next in the narrative (see Hale, 2016, for a recent review).

Early eye-tracking studies have shown that predictable words in sentences receive less fixations compared to less predictable words (Boston, Hale, Kliegl, Patil, & Vasishth, 2008; Demberg & Keller, 2008; Frank & Bod, 2011; McDonald & Shillcock, 2003; N. J. Smith & Levy, 2013, see Staub, 2015, for review). Recently, information-theoretic metrics have been tested against neuroimaging data to explore whether there is support for the idea that the brain might use statistics of the input implicitly to generate and evaluate predictions for efficient processing.

Work using fMRI has delineated the neural correlates of information-processing metrics in brain regions (see Bachrach, 2008, for early developments). In a study by Willems, Frank, Nijhof, Hagoort, and van den Bosch (2016), word entropy was shown to be negatively related to hemodynamic responses in the right inferior frontal gyrus, the left ventral premotor cortex, left middle frontal gyrus, supplementary motor area, and the left inferior parietal lobule, whereas word surprisal showed positive relationship bilaterally in the superior temporal lobes and in a set of (sub)cortical regions in the right hemisphere. Henderson, Choi, Lowder, and Ferreira (2016) have shown that syntactic surprisal is related to activity in the left inferior frontal gyrus and in the left anterior temporal lobes during natural story reading. Lopopolo, Frank, van den Bosch, and Willems (2017) further showed that fluctuations in part-of-speech, word, and phoneme perplexity during auditory narrative comprehension are related to largely separated brain activity in the temporal, inferior parietal and perisylvian cortical areas.

Electrophysiological methods are beginning to shed light on spectro-temporal neural correlates of probabilistic models. Word surprisal has been shown to correlate with the EEG N400 amplitude, a well-known marker of language processing, during sentence reading (Frank, Otten, Galli, & Vigliocco, 2015, see also Rabovsky & McRae, 2014; Rabovsky, Hansen, & McClelland, 2018, for work on simulating N400 amplitudes with analogues of surprise in connectionist models of reading). In a recent ECoG study, Nelson, Karoui, et al. (2017) showed that bigram word entropy during sentence reading is negatively related with fluctuations in the high gamma (70–150 Hz) power on electrodes overlying left posterior temporal regions. In a separate analysis of the same dataset, entropy reduction (changes in word-by-word syntactic uncertainty) was found to have a positive linear relationship with high gamma power in the left anterior and posterior inferior temporal electrode sites (Nelson, Dehaene, Pallier, & Hale, 2017).

Although fMRI work provided insights into spatial fingerprints of probabilistic computations and work in electrophysiology showed their temporal and high-frequency neural characteristics during sentence reading, it is as of yet unclear whether these statistical quantities are also reflected in ongoing electrophysiological activity during natural language listening. How are word-by-word context-dependent probabilistic predictions reflected in brain dynamics during naturalistic story comprehension?

To address this question, we recorded MEG during auditory story comprehension and used probabilistic language models to quantify the amount of information conveyed by each heard word (word perplexity) and the uncertainty as to what word(s) will likely follow up the current word (word entropy) (see figure 1, panel A). We then compared the power of frequency-specific MEG neural source dynamics as a function of high and low perplexity and entropy states (figure 1, panel B). We focused on dynamics in the theta, alpha, beta, and gamma frequency ranges as these have been previously implicated in core linguistic computations (see Bastiaansen & Hagoort, 2006; Lewis & Bastiaansen, 2015; Meyer, 2018; Weiss & Mueller, 2012, for reviews).

**Figure 1:**
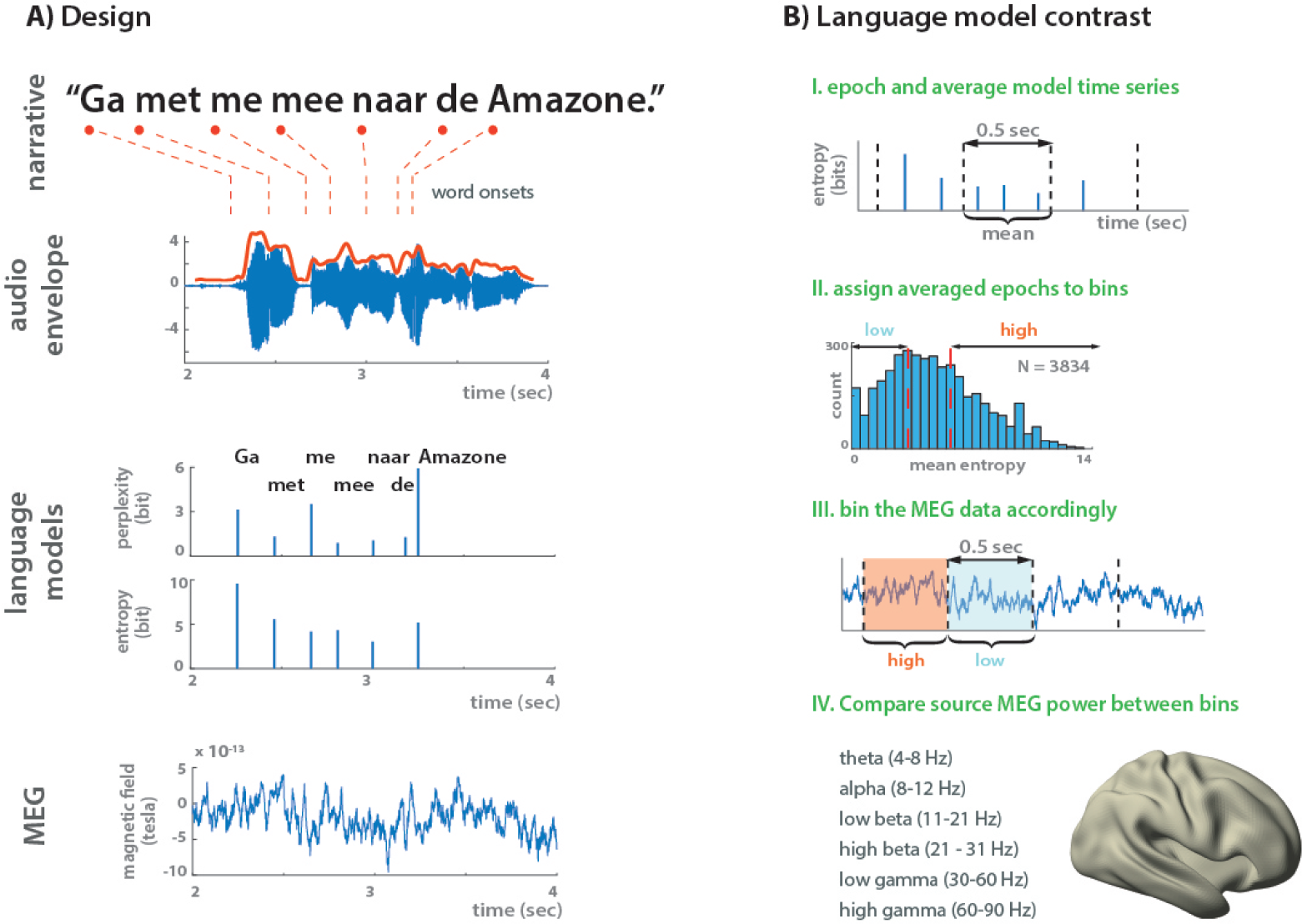
Design and data analysis. A) We recorded MEG responses while participants listened to auditory stories (excerpts from novels recorded as audiobooks). For each individual word in a story, we computed a complexity score (word perplexity or entropy) on the basis of trigram language model probabilities. B) Full story datasets were cut (epoched) into 0.5 second long snippets. On the basis of the average linguistic complexity within each snippet, we assigned the snippets into high or low complexity bins. We then compared frequency-specific power of MEG source activity across the complexity bins.

## 2 Methods

### 2.1 Participants and data acquisition

We recruited 25 healthy left- or right-handed (see Willems, Haegen, Fisher, & Francks, 2014, for inclusion justification) participants (13 female, mean age 25.8 *±* 7.6 [*M ± SD*]). Participants gave informed consent and were financially compensated for their participation. The study was approved by the “Committee on Research Involving Human Participants” (CMO) in the Arnhem—Nijmegen region and followed the guidelines of the Helsinki declaration. Participants received monetary compensation for the participation.

We collected MEG data with a 275 axial gradiometer system (CTF) in seated position at the Donders Centre for Cognitive Neuroimaging in Nijmegen, The Netherlands. The signals were digitized at a sampling frequency of 1200 Hz. Three coils were attached to the participant’s head (nasion, left, and right ear canals) to determine the position of the head relative to the MEG sensors. Throughout the measurement, the head position was continuously monitored using custom software (Stolk, Todorovic, Schoffelen, & Oostenveld, 2013). During breaks, the participant was allowed to reposition to the original position if needed. Participants were able to maintain a head position within 5 mm of their original position. Three bipolar Ag/AgCl electrode pairs were used to measure the horizontal and vertical electro-oculogram, and the electro-cardiogram.

### 2.2 Experimental procedure and stimulus materials

In the present analysis, we combined two sets of recordings of in total 25 participants (7 from the first set and 18 from the second set). In both datasets, participants listened to the same short stories in Dutch. The number of heard stories for the first ten subjects varied from 6 to 8. All subjects in the second dataset heard 5 stories. All subjects were instructed to listen to stories attentively for comprehension. The 18 subjects of the second dataset were additionally informed that they would answer two short multiple choice comprehension questions after each story (see Appendix A). They responded to comprehension questions by means of a button press.

Each story was presented binaurally via a sound pressure transducer through two plastic tubes terminating in plastic insert earpieces. A black screen was maintained while participants listened to the stories. Presentation of the auditory stories was controlled with Presentation software (version 16.4, NeuroBehavioral Systems Inc.). During story listening, participants were looking at a black screen and were not otherwise constrained.

The auditory stories were obtained from the subcorpus of Dutch literary stories available in the Spoken Dutch Corpus, “Corpus Gesproken Nederlands” (Oostdijk, 2000). The recordings were excerpts (mostly chapters) from audiobooks that were originally produced for Dutch Libraries for the Blind. The excerpts were spoken at a normal rate, in a quiet room, by different speakers (one speaker per story). As part of the Spoken Dutch Corpus project, word onset times and word offset times were determined (see Martens et al., 2002, for details), which we used for assigning language model output values to individual words (see section 2.3). The details for each of the stories are reported in Table 1.

**Table 1.** The number of words tokens, number of sentences, average sentence length (in number of words), and average word duration for each of the stimulus stories. Data source for Table 1: `table1.txt`. or `table1.Rdata`. Source code: `table1.R`.

| story | duration (mm:ss) | N. words | N. sen. | avg. sen. len | avg. word duration (sec) |
| --- | --- | --- | --- | --- | --- |
| 1 | 03:44 | 518 | 33 | 15.2 | 0.30 |
| 2 | 03:58 | 649 | 42 | 15.1 | 0.28 |
| 3 | 12:46 | 1730 | 131 | 13.1 | 0.29 |
| 4 | 03:59 | 657 | 35 | 18.2 | 0.27 |
| 5 | 07:55 | 1320 | 68 | 19.1 | 0.27 |
| 6 | 08:02 | 1127 | 87 | 12.8 | 0.28 |
| 7 | 04:03 | 549 | 34 | 15.7 | 0.28 |
| 8 | 08:04 | 999 | 62 | 15.9 | 0.29 |

### 2.3 N-gram language models

Word probabilities were estimated with a trigram language model, also known as a third order Markov model. A trigram language model relies on the simplifying (Markov) assumption that the probability of occurence of the current word depends on the past two words only, rather than on the entire preceeding string of words. The probability *P* of the of the word *w_t_* is thus conditioned on the past two words in the sentence:

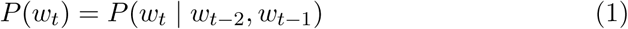

The stories were tokenized with the Frog natural language processing toolkit (van den Bosch, Busser, Canisius, & Daelemans, 2007). Language models for word entropy and perplexity estimates were then computed using the SRILM (Stolcke, 2002) and WOPR (van den Bosch & Berck, 2009) software packages, respectively. The model was trained on a selection of 10 million sentences (comprising 197 million word tokens; 2.1 million types) from the Dutch Corpus of Web, NLCOW (Schäfer & Bildhauer, 2012).

#### 2.3.1 Lexical perplexity

High word perplexity values indicate that the currently encountered word was less expected given the context. In psychological terms, perplexity is a “backward-looking” metric in that it models the degree of listener’s surprise upon encountering the word (higher perplexity means higher surprise) and the amount of information processing work required (higher surprise requires more information processing) given the past context. Perplexity is mathematically related to the information-theoretic measure of surprisal which is defined as the negative logarithm of the word’s conditional probability of occurrence:

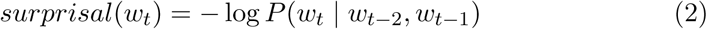

Perplexity is defined as the exponential transformation of surprisal:

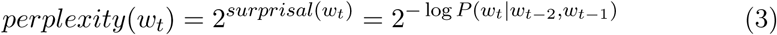

#### 2.3.2 Lexical entropy

Word entropy quantifies uncertainty in the probability distribution of possible upcoming words. High entropy signifies that there are many possible words that can follow the current word, whereas low entropy indicates that there are only few, highly probable words that can complete current sentence position. In other words, word entropy is a “forward-looking” metric and models the degree of the listener’s or reader’s uncertainty about the upcoming word given the words encountered so far. Entropy for the current word position is defined as the information-theoretic chaos in the distribution of all possible upcoming words at *t* + 1 given the words encountered so far (*w*_1_*_,…,t_*):

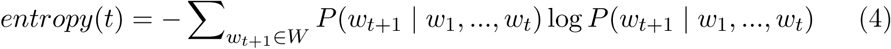

where *W* denotes the set of possible words following (*w*_1_*_,…,t_*).

We used base-2 logarithm as the scaling factor, therefore both perplexity and entropy are expressed in bits. A descriptive visualization for word perplexity and word entropy along with two control variables (lexical frequency and word duration) is provided in figure 2. Along the diagonal, the matrix shows distributions for word entropy, perplexity, lexical frequency, and word duration (msec) per story. Distributions for lexical perplexity and lexical frequency have been log 10-rescaled for visualization purposes. Lower off-diagonal panels display scatterplots for combinations of variables. The corresponding upper off-diagonal panels show the corresponding Pearson correlation coefficient between variables.

**Figure 2:**
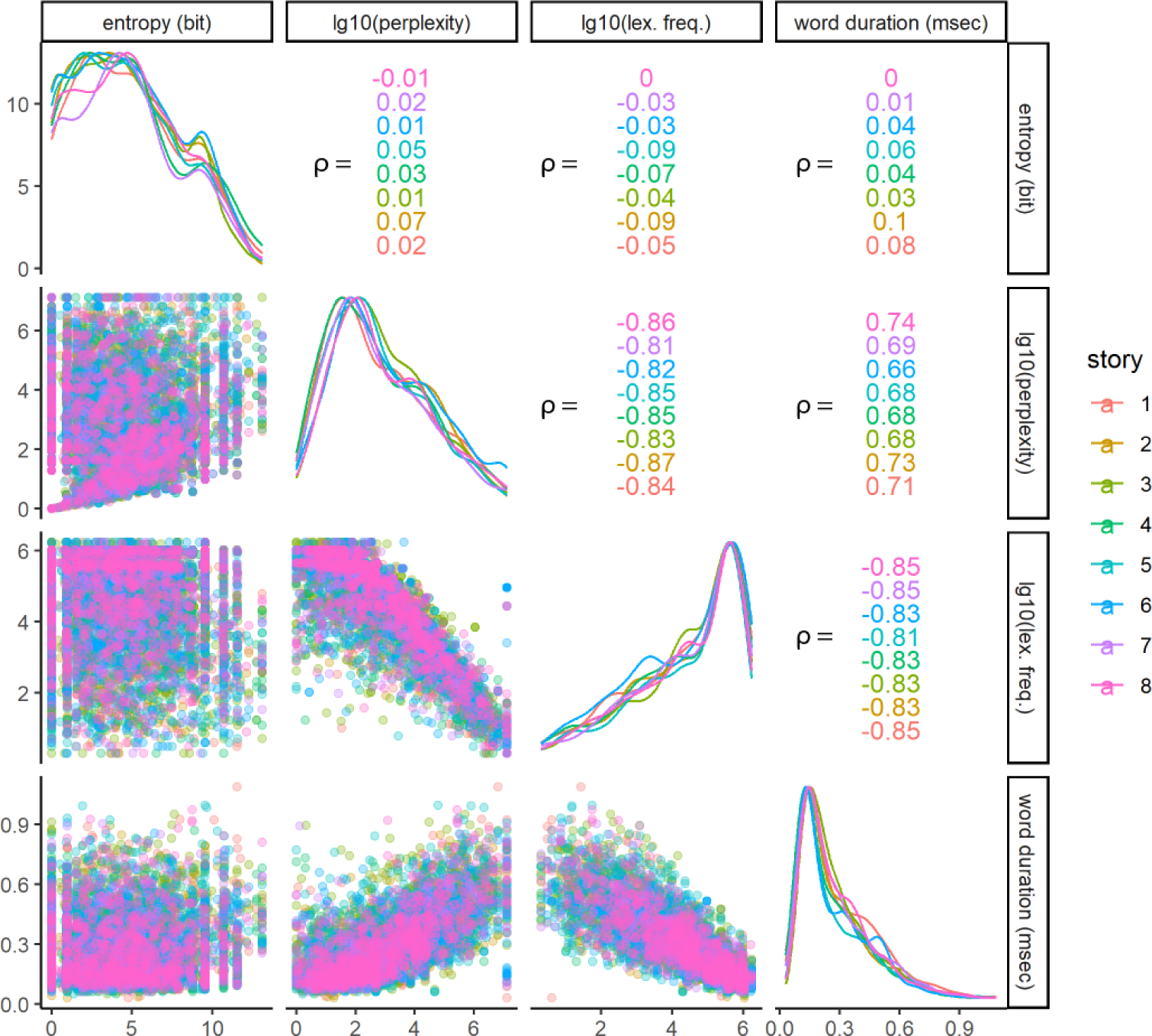
Scatterplot matrix for entropy, log10-transformed perplexity, log10-transformed lexical frequency, and word duration (msec) per story. Color-coding represents data for individual stories. Values in the upper off-diagonal panels are story-specific Pearson correlation coefficients for the corresponding variable pair. On the diagonal are story-specific density plots for the corresponding variable. Lower off-diagonal panels show scatterplots for the corresponding variable pair. Data source: `language_data.txt, stimuli_table.txt`, R code to reproduce the scatterplot matrix: `figure_2.R`

### 2.4 Preprocessing

The analysis of MEG data was performed in MATLAB, version 2016b (Mathworks, Inc.) with the FieldTrip software package (Oostenveld, Fries, Maris, & Schoffelen, 2011), together with a custom in-house MATLAB code. The complete code used in the analysis with cursory documentation is available at: https://github.com/KristijanArmeni/dyncon_streams/tree/master.

#### 2.4.1 MEG

Prior to preprocessing, the raw data were demeaned. We then applied notch filtering at the bandwidth of 49–51, 99–101, and 149–151 Hz to remove the potential line noise artifacts. Artifacts related to muscle contraction and squidjumps were identified and removed using a semi-automatic artifact rejection procedure (http://www.fieldtriptoolbox.org/tutorial/automatic_artifact_rejection). The data were then downsampled to 300 Hz. MEG components reflecting eye-blinks were estimated using the FastICA algorithm (https://research.ics.aalto.fi/ica/fastica/) as implemented in Fieldtrip functionalities
. Relevant components were identified based on their topography and time-courses and removed from the data.

#### 2.4.2 MRI

Original volumes of T1-weighted MRI images of each participant were manually co-registered to the MNI headspace coordinate system. MRI images were then re-sliced to an isomorphic 256 × 256 × 256 space and co-registered to the MEG (CTF) headspace coordinate system by defining the positions of pre-auricular points and the nasion MEG coil on the re-sliced MRI images. To reproduce and localize the position of the left and right pre-auricular points on the MRI images, we placed custom-made markers (vitamin E capsules) in participants’ ear molds during MR image acquisition (see http://www.fieldtriptoolbox.org/faq/how_are_the_lpa_and_rpa_points_defined). To make the co-registration as precise as possible, we used the same ear molds as in the MEG session. No specific marker, other than anatomical identification on the image, was used to localize the nasion coil.

After co-registration, the Brain Extraction Tool (BET; S. M. Smith, 2002) from the FSL command-line library (v5.0.9; Jenkinson, Beckmann, Behrens, Woolrich, & Smith, 2012) was used to delete the non-brain tissue (skull striping) from the whole head. To obtain a description of individual subject’s cortical sheet, we performed cortical surface reconstruction with the Freesurfer image analysis suite, which is documented and freely available for download online (http://surfer.nmr.mgh.harvard.edu/), using the surface-based stream implemented in the recon_all command-line tool. The post-processing of the reconstructed cortical surfaces was performed using the Connectome Workbench wb_command command-line tools (v1.1.1; https://www.humanconnectome.org/software/workbench-command).

#### 2.4.3 Audio

The onsets of the acoustic signals from audio stories were first corrected for the delay with respect to the MEG triggers. We then computed the amplitude envelope of auditory signals following Gross et al. (2013). Using the Chimera toolbox (http://research.meei.harvard.edu/chimera/More.html, see also Z. M. Smith, Delgutte, & Oxenham, 2002), we constructed 10 frequency bands of equal width in the range 100–10,000 Hz to match the frequency range of human hearing along the basilar membrane. The power envelopes of each of the constructed frequency bands were averaged together to yield a single auditory power envelope. This averaged auditory envelope was used in computation of audio–MEG coherence (see Section 2.7.3) and as confounding variable in the analysis of linguistic variables (see Section 2.7.2).

### 2.5 Epoching and linguistic information content

The pre-processed MEG data were re-epoched into 0.5 second-long time segments (see figure 1, panel B). To allow for spectral estimates that are both robust and qualitatively comparable across individual epochs, we decided to keep the epoch-length (i.e. number of samples used in estimation) fixed. Aligned with recorded MEG time series, we logged the timing of word onsets and offsets and the word’s entropy and perplexity values (see Section 2.2 for how onsets were determined). Since an individual epoch could straddle several words (our epoching step did not consider word boundaries, but see supplementary information for an additional analysis which did), we computed, for each epoch, the mean over individual word perplexity and entropy values weighted by the individual word’s duration (such that longer words contributed more to the overall mean complexity score). This step resulted in a point estimate of linguistic complexity in each epoch.

The distribution of all epoch-specific complexity values was subsequently binned into low and high complexity bins which contained entropy or perplexity scores below the 33- and above the 66-quantiles, respectively (see figure 1, panel B, third sub-figure). The two linguistic complexity bins served as the contrast for our comparison of frequency-specific MEG source power (see section 2.8).

### 2.6 Temporal lagging of MEG time-series

An important caveat of the procedure described so far is that, despite temporal smoothing due to the averaging step per epoch, it assumes quasi-instantaneous effects of linguistic features on MEG dynamics (e.g. a word’s unexpectedness is assumed to be observed at time of word onset in MEG time-series). To adjust for this, we repeated the epoching procedure 3 times, each time with temporally lagged versions of MEG signals relative to the time course of linguistic information. That is, prior to epoching, we selected the MEG time-series starting 200, 400, and 600 msec post-onset and realigned them with the onset of the original language time-series.

Temporal shifting of the estimated MEG power relative to the linguistic features allowed us to investigate the temporal dynamics of the association between oscillatory power and the linguistic features, accounting for a likely delay between the word onset and the modulatory effect of the linguistic feature on the brain response. However, we should stress that our choice of epoching and averaging complexity metrics described in the preceding section in principle does not allow us to make *strong* statements that link the observed effects to linguistic features of individual words; but rather that any observed effects reflect slow fluctuations of the linguistic features aggregated across several words.

### 2.7 Source reconstruction

The cortical sheet reconstruction procedure (see section 2.4.2) resulted in a description of individual subjects’ locations of potential neural sources along the cortical sheet (source model) with 7,842 source locations per hemisphere. We used a single-shell spherical volume conduction model (head model) based on a realistic shaped surface of the inside of the skull (Nolte, 2003) to compute the forward projection matrices (leadfields).

We performed “dynamic imaging of coherent sources” (Gross et al., 2001), a frequency-domain spatial filtering technique, to estimate the frequency-specific single-trial audio-MEG coherence and MEG neural source power on the reconstructed cortical sheets. To estimate frequency-resolved coherence spectra we used linearly constrained minimum variance spatial filtering (LCMV, Veen, Drongelen, Yuchtman, & Suzuki, 1997). Both methods can be deployed with the Fieldtrip “ft_sourceanalysis” routine.

#### 2.7.1 Neural source power

The cross-spectral density of the sensor-level data was first computed by estimating the single-trial Fourier power per frequency of interest. The dpss multitaper method with progressively larger smoothing windows (2 Hz for theta and alpha bands; 5 for beta bands; and 15 for gamma bands) was used to estimate the power at the following frequency ranges: theta (4–8 Hz; centered at 6 Hz), alpha (8–12 Hz; centered at 10 Hz), low beta (11–21 Hz; centered at 16 Hz), high beta (21–31 Hz; centered at 26 Hz), low gamma (30–60 Hz; centered at 45 Hz), and high gamma (60–90 Hz; centered at 75 Hz) frequency bands. The Fourier spectra were subsequently converted to bi-variate cross-spectral densities.

The subject-specific leadfields (see section 2.4.2) and cross-spectral densities were then used to estimate the inverse spatial filters (beamformers). In order to reduce the sensitivity to noise and to increase the consistency of the spatial maps across subjects, the lambda regularization parameter was specified as 100%. This step resulted in a spatial filter for each source location on the cortical sheet. The sensor level single-trial (MEG channel-by-trial) Fourier-transformed power data were then left-multiplied with the location-specific spatial filter (source dipole moment-by-MEG channel) to yield the trial-specific estimate of source power at that location.

#### 2.7.2 Confounding variables

Initial exploration of our independent variables shows that our trigram word perplexity is negatively related with word lexical frequency (see figure 2). That is, the model assigns higher perplexity scores on average to words that occur less frequently in the corpora. We therefore decided to treat mean log-transformed lexical frequency per epoch as a potential confounding variable in our analysis. Word frequency counts were obtained from the SUBTLEX corpus (Keuleers, Brysbaert, & New, 2010).

In addition, to remove the part of the variance in the estimated MEG oscillatory activity that is due to purely low-level acoustic fluctuations in the input signal, we included epoch-averaged auditory envelope power (see section 2.4.3) as a potential confound as well. Prior to computing the statistical contrast described in section 2.8, the variance attributed to lexical frequency and auditory envelope fluctuations was estimated by means of a general linear model and was regressed out from the power spectra (see Stolk et al., 2013, for details).

#### 2.7.3 Audio envelope–MEG source coherence

To estimate the coherence between averaged audio envelope and source neural dynamics, we first computed a complex Fourier representation of the sensor level signals per epoch centered at 6 Hz (*±* 2 Hz). The Fourier spectra were subsequently converted to bi-variate cross-spectral densities. The subject-specific leadfields (see section 2.4.2) and cross-spectral densities were then used to estimate the inverse spatial filters (beamformers). In order to reduce the sensitivity to noise and to increase the consistency of the spatial maps across subjects, the lambda regularization parameter was specified at 100%, which reflects a diagonal loading of the co-variance matrix with the mean of the variance across channels.

#### 2.7.4 Frequency-resolved audio envelope–MEG source coherence

To estimate audio-cortico coherence across a broader spectrum of frequencies, we used the LCMV approach, a time-domain beamformer algorithm. The story-epoched MEG time-series were first re-epoched to 4s-long time windows. We then computed the sensor-level co-variance matrix that was used in the LCMV algorithm to estimate the beamformer weights. The lambda regularization parameter to “ft_sourceanalysis” routine was specified at 100%.

We only computed spatial filters for brain parcels that showed maximal theta coherence in the DICS source reconstruction (Section 2.7.3). Parcellation (grouping of source points into brain areas or parcel) was defined with the Conte69 atlas (brainvis.wustl.edu/wiki/index.php//Caret:Atlases/Conte69_Atlas), which provides a parcellation of the neocortical surface based on Brodmann’s cytoarchitectonic atlas, consisting of 41 labeled parcels per hemisphere. The sensor-level time-series were then projected to the source space by left-multiplying them with location-specific beamformer weights.

Using the dpss multitaper method with a smoothing window of 1 Hz, we computed the complex Fourier representation of the time-domain audio and MEG source signals in the range between 0 and 50 Hz which was used to compute the coherence spectrum per frequency band.

### 2.8 Within subject contrast

An independent two-samples *t*-statistic was used to quantify differences in mean neural source power between high and low complexity levels per each of the six frequency bands in every subject. The *t*-statistic was used in order to normalize for potential signal-to-noise differences in MEG signals between the two groups not related to the comparison of interest. This step resulted in a *t*-map (a number-of-source-locations by number-of-time-lags matrix of *t*-values) for every subject and frequency band.

### 2.9 Group contrast and inference

The subject-specific *t*-statistics computed in the first-level step were entered into a group-level analysis. The dependent-samples *t*-statistic was computed for every pair of source-location and time-point where the first sample consisted of observed per-subject independent-samples *t*-statistics from the first-level analysis, and the second sample consisted of a matrix of same dimensions than that sample 1 but filled with zeros instead of observed *t*-statistics.

Statistical significance of observed group differences was determined by means of the cluster-based non-parametric permutation test (Maris, Schoffelen, & Fries, 2007). Here, the goal is to determine whether the global null hypothesis can be rejected (i.e. that there is at least one cluster across all source, time, and frequency domains which makes data non-exchangeable between experimental conditions). For each frequency-band, we included temporally and spatially adjacent sample-specific *t*-statistics that exceeded the alpha level of 0.05 into clusters.

For each detected cluster, a single *cluster statistic* was calculated by summing the sample-specific *t*-statistics from that cluster. Next we computed a reference (null) permutation distribution to which we compared our observed cluster statistics. The reference distribution was created by permuting (randomly exchanging) data between the conditions, and then calculating the maximal positive and negative cluster statistics for each permuted data set. The permutation step was repeated 1,000 times. This step hence resulted in two distributions of 1,000 largest and smallest summed cluster statistics for each of the six frequency-bands, **D***^pos^* and **D***^neg^*,ℝ ∈ 6 × 1000.

Finally, to reject or accept the global null hypothesis (i.e. the hypothesis stating that the data are randomly exchangeable between high and low conditions in space, time, and frequency domains), we created the final permutation distribution of cluster statistics by taking the largest and the smallest summary statistic across the six frequency bands. This step resulted in two distributions, **d***^pos^* and **d***^neg^*, ℝ ∈ 1 × 1000. We then selected the largest and the smallest summary statistics from the distribution of observed summary statistics. Finally, we computed the proportion of permutation statistics (in **d***^pos^* and **d***^neg^*) that are greater (smaller) or equal to the observed largest (smallest) cluster statistic (*p*-value). The global null hypothesis was rejected if the proportion was smaller than 2.5 % (*α*-level). This step resulted in two *p*-values, one for each direction. The smaller *p*-value of the two was taken as the result (and direction) of the statistical test (i.e. this *p*-value is reported in the results section). The permutation test controls for multiple comparisons in space, time, and frequency.

## 3 Results

### 3.1 Comprehension questionnaires

The subjects who had to answer the comprehension questions (see Section 2.2) all correctly responded to at least 7 or more (out of 10) comprehension questions showing that they did pay attention to the contents of the narratives.

### 3.2 Language model contrasts

We compared the band-limited MEG signal power in high and low entropy and perplexity bins (see section 2.5) across 6 frequency bands (theta, alpha, low-beta, high-beta, low-gamma, high-gamma) and 4 time points (0, 200, 400, 600 msec).

#### 3.2.1 Word perplexity

Comparison across frequencies showed that there was a statistically significant difference in MEG power between high and low perplexity epochs (permutation *p*-value = 0.018, negative direction). For interpretation of the global effect, we determined in which frequency bands power comparison showed the largest differences and thus likely driving our global effect. To do so, we used a heuristic where we inspected which frequency-specific permutation *p*-values were smaller than *α* = 0.025. The smallest *p*-values were observed for the low and high beta-bands (clusters displayed in figure 3.2.1) meaning that, on average, highly surprising words in the narrative were accompanied by reduced beta-band power relative to expected, unsurprising words.

**Figure 3:**
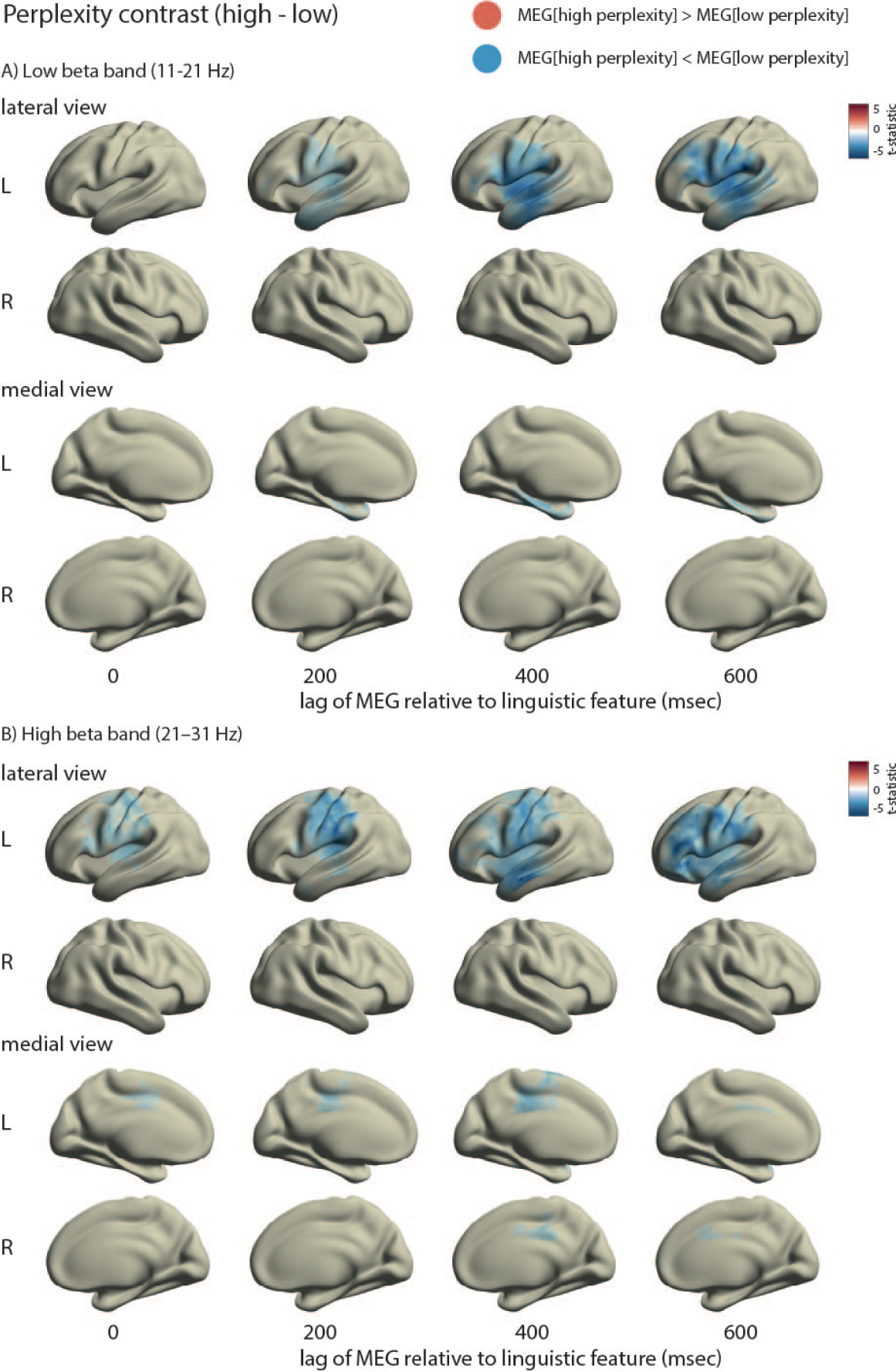
Group level *t*-maps for low beta (11–21 Hz, panel A and high beta band (21–31 Hz, panel B) power quantifying differences between high and low perplexity bins. Source maps show paired-samples *t*-statistic per source location and time lag where the upper and lower extreme point of the color bar are determined by the maximal absolute value over *t*-statistics and its negative, respectively. For the purposes of visualization, displayed are only *t*-statistics for source locations belonging to the permutation cluster with the highest cluster statistic, that is, source locations not belonging to the maximal clusters the cluster are set to a value of zero. MATLAB code to reproduce source maps: `script_sourceplot4eps.m`

For both frequency bands, the effects are predominantly left-lateralized with peaks over left central, temporal and frontal areas. Power differences progressively increase with each subsequent temporal lag peaking at the lags of 400 and 600 msec.

#### 3.2.2 Word entropy

Comparison between high and low entropy epochs showed a trend towards significance (permutation *p*-value = 0.033, positive direction). Frequency-specific clusters with *p*-values exceeding the *alpha* threshold of 0.025 were observed the theta (positive cluster) and low-beta (negative cluster) bands (clusters displayed in figure 4). Hence, the global entropy effect was likely driven by differences in the theta and beta bands.

**Figure 4:**
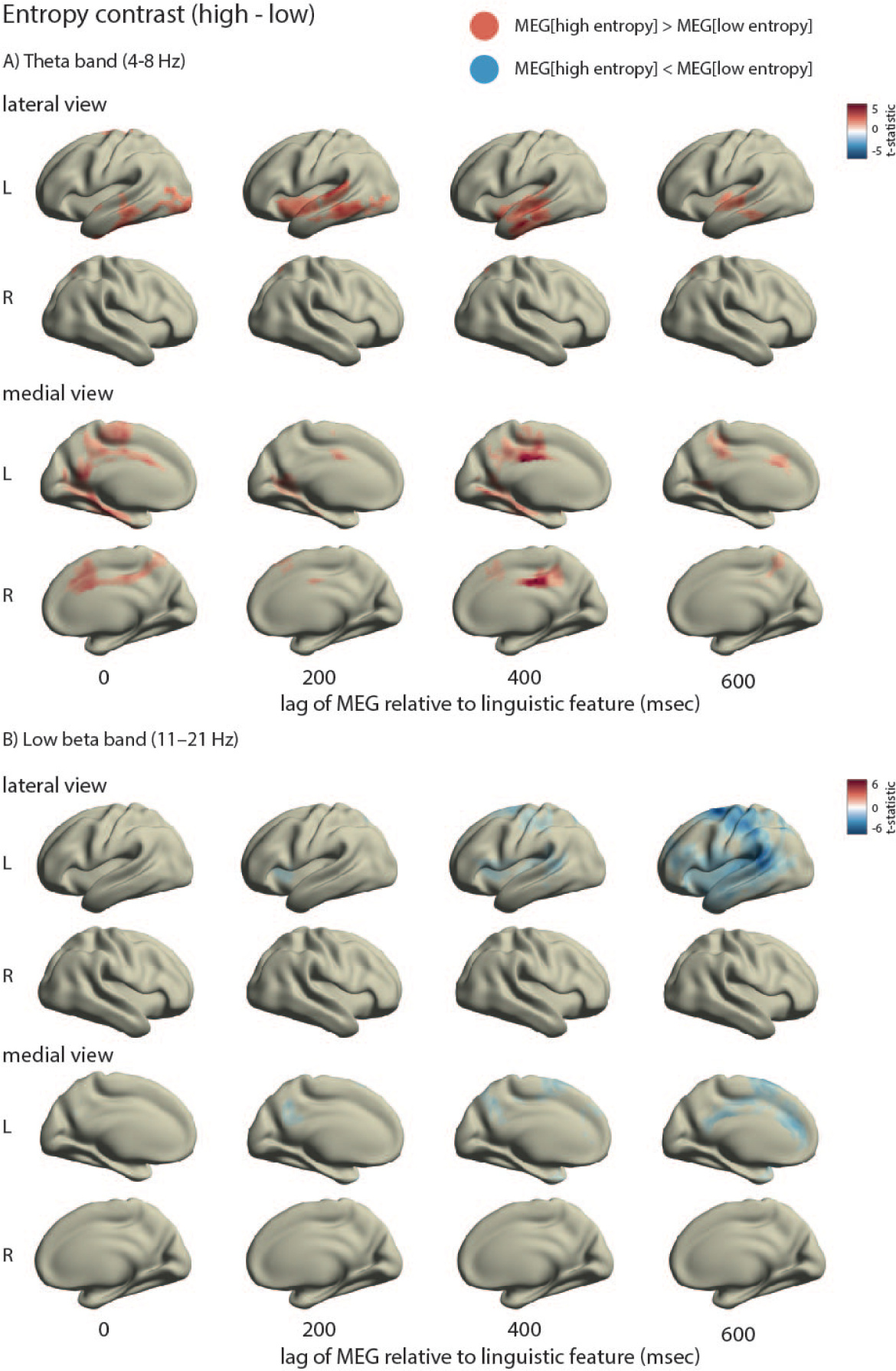
Frequency-specific power differences between conditions visualized here as group-level *t*-maps for theta-band (4–8 Hz, panel A) and lower beta-band (11–21 Hz, panel B) power quantifying differences between high and low entropy bins. Source maps show paired-samples *t*-statistic per source location and time lag where the upper and lower extreme point of the color bar are determined by the maximal absolute value over *t*-statistics and its negative, respectively. For the purposes of visualization, displayed are only *t*-statistics for source locations belonging to the permutation cluster with the highest cluster statistic, that is, source locations not belonging to the maximal clusters the cluster are set to a value of zero. MATLAB code to reproduce source maps: `script_sourceplot4eps.m`

The low beta-bands revealed predominantly negative differences (blue color coding in figure 4, panel B) meaning that in high entropy contexts beta-band power was on average reduced relative to more predictive, low entropy contexts. The spatial structure of observed group-level beta-band differences across time lags shows a predominantly left-lateralized peak differences over left central areas and superior temporal areas extending to angular gyrus. Inspecting the temporal structure of the effect, differences are most pronounced when the MEG signal is lagged relative to the linguistic feature by 600 msec.

Power differences in the theta-band showed a predominantly positive direction (red color coding in figure 4, panel A) showing that high entropy contexts, which are less predictive about upcoming words, are on average accompanied by increased theta-band power compared to more predictive, low entropy contexts.

Theta-band power power differences showed left-lateralized differences with focal maxima over the middle temporal and inferior frontal lobes (red color coding in figure 4, panel A, upper). We additionally localized differences to medial cortical areas with focal peaks in the bilateral posterior cingulate cortex and posterior medial temporal areas, right anterior cingulate cortex, and left inferior parietal cortex. (figure 4, panel A, lower).

### 3.3 Audio envelope–MEG coherence

To additionally ascertain that our entropy-theta trend is not driven by the relationship between acoustic envelope fluctuations and MEG power spectrum we, as a control analysis, explored the topography of this effect in our dataset. This phenomenon, also known as “speech entrainment”, has been extensively reported in studies using auditory sentences or narratives as stimuli (Ahissar et al., 2001; Giordano et al., 2017; Gross et al., 2013; Lam, Hultén, Hagoort, & Schoffelen, 2018; H. Luo & Poeppel, 2007; Park, Ince, Schyns, Thut, & Gross, 2015).

We computed coherence between the source reconstructed MEG time-series and speech envelope fluctuations. The spatial distribution of average audio envelope-MEG coherence for the theta band is displayed in figure 5, panel A. Panel B shows the audio envelope-MEG coherence spectra (individual participants in grey) for those source parcels that showed maximal dics coherence values in the corresponding hemisphere.

**Figure 5:**
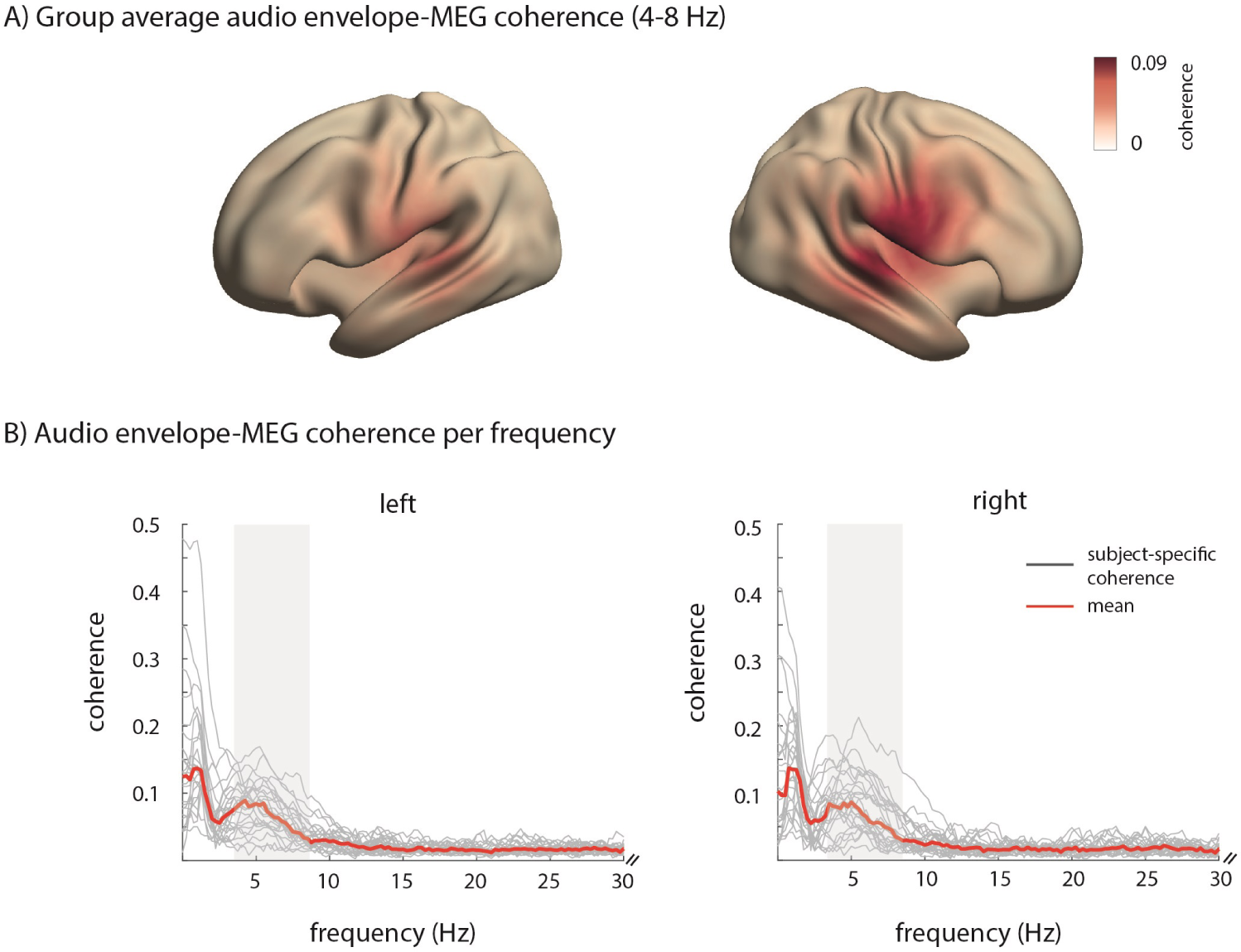
Audio envelope–MEG coherence quantifying entrainment of low-frequency MEG power by auditory envelope fluctuations. A) Coherence between auditory envelope and MEG signals in the theta band. Visualized are coherence values averaged across subjects. Coherence is bounded between 0 and 1. B) Coherence between auditory envelope and MEG signals across the frequency spectrum in the 1–50 Hz range (shown up to 30 Hz) in parcels with maximal coherence across the corresponding hemisphere. Grey lines show audio-MEG coherence per subject, red line shows the mean across subjects. Shaded areas mark the frequency range represented in the source plots in panel A. Code to reproduce source maps: `script_fig5.m`

## 4 Discussion

In the current study, we tested how frequency-specific oscillatory brain dynamics relate to expectation-based predictive processing in naturalistic language comprehension. We used well-defined information-theoretic metrics as an objective measure for how unexpected each heard word was given the past two words (word perplexity) and how predictable the upcoming input was (word entropy). Our approach allowed us to study brain dynamics in response to naturalistic, acoustically presented, linguistic signals. We observed predominantly left-lateralized effects, with low and high beta-band dynamics relating to fluctuations in perplexity and entropy, and theta-band dynamics to entropy modulations.

### 4.1 Beta-band dynamics reflect maintenance of context

Comparison of unexpected, high-perplexity words with expected, low-perplexity words showed that unexpected words were accompanied by lower neural source power in both the low and high beta-bands. Put differently, more expected words led to higher beta-band power. Beta-band dynamics were also negatively related to stimulus entropy, that is, lower beta-band MEG source power was observed for high entropy (less predictive) contexts relative to low entropy (more predictive) context.

The beta effects reported are in line with the accounts proposing that decreases in beta-band power signal new processing demands in the neural system (Engel & Fries, 2010; Lewis & Bastiaansen, 2015; Weiss & Mueller, 2012). In the present study, a decrease in beta-band power upon encountering unexpected words possibly signals the need to revise and update the current representation of sentence and story contents. In contrast, if the words are less perplexing given the context, no substantial additional neural processing is needed to update the current contextual state, and beta-band dynamics are unperturbed. Similarly with entropy; in more predictable contexts, strong predictions are licensed which leads to sustained beta-band dynamics in higher-level areas, possibly providing input to areas involved in lexical or sensory processing.

The left-lateralized topography of perplexity-driven and entropy-driven beta-band dynamics over central, temporal, and frontal areas supports its hypothe-sized general role of reflecting maintenance of contextual states of the system for top-down predictions (Lewis & Bastiaansen, 2015) and is in line with previous findings from sentence reading studies reporting left frontal beta-band reduction to semantic and syntactic violations in sentences (Bastiaansen, Magyari, & Hagoort, 2010; Kielar, Panamsky, Links, & Meltzer, 2015; Wang, Jensen, et al., 2012), and left frontal sensor-level beta reductions to incongrouent discourse-level information in short paragraphs (Lewis, Schoffelen, Hoffmann, Bastiaansen, & Schriefers, 2017).

Recent research has also linked beta-band dynamics with sources in sensorimotor areas to the generation of temporal predictions to the auditory brain areas during auditory perception (Morillon & Baillet, 2017). Further, beta-band dynamics in left frontal areas have been found to modulate the strength of speech-brain coupling in auditory cortical areas during auditory story comprehension (Keitel, Ince, Gross, & Kayser, 2017). The exact computational role of (pre)motor and frontal areas in online speech perception and language comprehension remains an ongoing area of research (Lima, Krishnan, & Scott, 2016; Skipper, Devlin, & Lametti, 2017) and is beyond the focus of the present study. Suffice it to note presently that the topography of beta effects substantiates the interpretation that beta-band dynamics are involved in maintaining active states of the system, possibly for top-down predictions (see Meyer, 2018; Spitzer & Haegens, 2017, for recent reviews).

The observed left-lateralized beta-band dynamics are most prominent when the MEG signal is lagged relative to linguistic information between 400–600 msec, which reflects the fact that the computations of prediction (dis)confirmation or context-driven uncertainty cannot happen instantly, upon hearing the current word, but is rather only possible once the words that follow the current word have been observed. Given that average word duration in our stimuli was approximately between 200 and 300 milliseconds, a lag of 400–600 milliseconds seems a plausible latency at which contextual engagement and comparison of predictions based on the current context can be observed in brain signals. However, as mentioned in section 2.6, our estimates of the temporal profile of these affects are tentative given temporal aggregation inherent in our analysis pipeline. Over all, these dynamics are in line with previous studies reporting beta-band reductions to semantically or syntactically incongruent (and hence, likely unexpected) words in sentences in late time windows spanning the broader time window of 0.5 to 1.5 sec post word onset (Bastiaansen et al., 2010; Kielar et al., 2015; Y. Luo, Zhang, Feng, & Zhou, 2010; Wang, Jensen, et al., 2012).

It is interesting to note that perplexity-driven beta dynamics showed an earlier onset relative to entropy-driven beta. This is not unexpected, given the fact that computation of word perplexity depends crucially on the actually observed word, whereas word entropy, strictly speaking, is not the property of any word alone, but is instead a function of the distribution of *possible upcoming words* at the current word position in a sentence, a computation that likely requires longer latencies for it to be reflected in cortical signals. Our results on temporal progression of spectral modulations are in line with earlier reports showing, among other, that lexical language model complexity metrics in general do not relate to early (peaking *<* 300 ms post word onset) language-related ERP components (Frank et al., 2015).

### 4.2 Theta dynamics relate to lexical uncertainty

In the current study, word entropy states show a trend towards a positive relationship with theta-band dynamics; that is, brain activity tends to be higher in high entropy (less predictable) contexts. Entropy quantifies the uncertainty about the possible upcoming words at the currently heard word. Entropy is high if the two-word context up to the current word position can be completed by many possible words and low in cases when there are only a few, highly probable continuations.

From a psycholinguistic perspective, higher oscillatory activity for unpredictable continuations relative to predictable ones can be interpreted as support for accounts emphasizing that the amount of neural information-processing resources used increases with increased “competition” (i.e. increased entropy) between plausible continuations (see Marslen-Wilson, 1987, for such an account in auditory word recognition). The present results therefore do not corroborate—at least in terms of theta-band oscillatory activity—opposite accounts postulating that use of predictive neural resources would be delayed until the commitment to specific predictions is possible, that is, more neural resources will be used (and more rapidly) in situations of low entropy (see Ettinger, Linzen, & Marantz, 2014, for such an account and evidence in morphology). Theta-band dynamics, as reported here, then likely reflect input-driven lexical computations as opposed to genuinely predictive computations *per se*.

Traditionally, theta band increases have been reported for semantically incongruent words relative to congruent counterparts (Bastiaansen & Hagoort, 2015; Y. Luo et al., 2010; Wang, Zhu, & Bastiaansen, 2012; Willems, Oostenveld, & Hagoort, 2008) and for unexpected words regardless of contextual predictability (Rommers, Dickson, Norton, Wlotko, & Federmeier, 2017). Such word-related theta-band modulations have been interpreted to reflect to long-term memory retrieval processes where less expected or incongruent words would be harder to retrieve from memory (see Bastiaansen & Hagoort, 2006; Meyer, 2018, for reviews).

On the one hand, our perplexity-based contrast (perplexity is a formal analogue of experimentally-derived word probabilities, see N. J. Smith & Levy, 2011, for discussion) did not reveal modulations in theta band as would be expected from the above-mentioned studies, hence our data do not directly support the theta-memory account. On the other hand, high entropy story contexts, with low degree of predictability (i.e. high degree of lexical competition between upcoming words), can be thought of as requiring more “effort” in the neural system to retrieve the correct word form and hence would be expected to show increased theta-band activity. This is indeed the pattern reported here. Thus, our results, despite not showing surprise-related theta increases can be subsumed under the aforementioned lexical competition (Marslen-Wilson, 1987) or related memory retrieval accounts (Meyer, 2018).

However, in addition to linguistically-motivated accounts, the present patterns of results can also be accounted for in the framework of probabilistic inference (Chater & Manning, 2006; Kuperberg & Jaeger, 2016; Pouget et al., 2013) where it is postulated that the listener is constantly weighing the relevance of the incoming lexical evidence (currently processed word) and prior knowledge (here the minimal sentential context so far) for updating and inferring the model of the world (narrative contents).

In high-entropy situations, when the sentential context is not strongly predictive of upcoming words, the corresponding state of the relevant neural circuits is less determined by their contextual history, but is instead more sensitive to the actually observed lexico-statistical properties of the incoming words (see Strange, Duggins, Penny, Dolan, & Friston, 2005, for a related interpretation of hippocampal modulations to stimulus entropy). Put differently, because the context affords only weak predictions, more information-processing work needs to be done on the currently observed words to foster evidence for inference and future efficient processing. Assuming that increases in theta-band MEG power reflect increased synchronous engagement of the underlying neuronal populations, the probabilistic processing account is corroborated by the fact that theta-band neural source power is higher in less vs. more predictive contexts.

Localization of the theta-band differences shows that these are strongly left-lateralized with a focal maximum in middle temporal areas, which adds credence to our interpretation that theta-band dynamics reflect increased sensitivity of neural systems to lexico-semantic evidence. In a recent meta-analysis, the left lateral temporal cortex has been suggested to be sensitive to lexico-semantic processing demands during sentence comprehension (Hagoort & Indefrey, 2014). In a recent MEG study, Lam, Schoffelen, Uddén, Hultén, and Hagoort (2016) source-localized word-related theta-band power decreases to left anterior temporal lobes. In addition, modulations in the theta-band have been consistently linked to word-related computations (i.e. lexical retrieval or lexical error computation) in sentence processing tasks (Bastiaansen & Hagoort, 2015; Bastiaansen, Linden, Keurs, Dijkstra, & Hagoort, 2005; Bastiaansen, van Berkum, & Hagoort, 2002; Hald, Bastiaansen, & Hagoort, 2006; Kielar et al., 2015; Wang, Zhu, & Bastiaansen, 2012).

Alongside theta differences in the left lateral temporal cortical areas, we also observed prominent peaks in several medial cortical structures. Activation of these regions is well documented in fMRI studies of pragmatic text comprehension beyond single sentences (Ferstl, Neumann, Bogler, & von Cramon, 2008). In the memory literature, these areas are considered as core components of the extended posterior medial memory system (Ranganath & Ritchey, 2012) which is thought, among others, to underlie our ability to maintain the construction of a mental representation of relations between entities denoted by the text, so-called situation models (Zwaan & Radvansky, 1998). Increased theta-band activity with high entropy could therefore be taken to signify increased use of neural resources for pragmatic and contextual computations in order to overcome high levels of lexical uncertainty induced by the text so far.

Timings of theta-band patterns are consistent with known temporal characteristics of event-related electrophysiological responses (but see our remark on temporal estimates in section 2.6). The N400 response has been consistently linked to (predictive) aspects of lexical processing (Kutas & Federmeier, 2011, see also Nieuwland et al., 2018, for a criticial replication attempt). In sentence reading, both and N400 effect and a spectral theta-band increase have been observed to unexpected words relative to expected ones (Rommers et al., 2017) suggesting that the two data analysis representations might reflect partially similar underlying biophysical processes.

Finally, a potential concern could be that our spectral and spatial entropy patterns in fact coincide with the so-called acoustic entrainment effects which are typically observed in the theta frequency range over left and right superior temporal cortex (Lam et al., 2018). However, our MEG-audio coherence analysis indicates that the underlying neural sources have distinct spatial profiles with modest or no focal overlap; that is, entrainment due to acoustic properties of the input shows sources with peaks in the superior temporal which are not left-lateralized areas as opposed to entropy-related patterns which are predominantly left-lateralized with peaks in the middle temporal lobes. This suggests that entropy-related theta band dynamics reflect electrophysiological processes beyond responses to acoustic-sensory inputs alone.

### 4.3 Comparison with related prior work

To the best of our knowledge, only three other studies so far explored the relationship between probabilistic language complexity in electrophysiology (albeit using sentence reading task rather than auditory narrative comprehension). Frank et al. (2015) reported no significant relationship between next-word entropy and language-related ERPs in sentence reading after accounting for the effects of surprisal. In a recent ECoG study, Nelson, Karoui, et al. (2017) showed that bigram word entropy during sentence reading is negatively related with fluctuations in high gamma (70–150 Hz) power on electrodes surrounding left posterior temporal regions. In a separate analysis of the same dataset, Nelson, Dehaene, et al. (2017) showed that sentential entropy reduction (modeling per-word changes in grammatical uncertainty) was reported to have a positive linear relationship with high gamma power in the left anterior and posterior inferior temporal electrode sites.

There are several differences between the studies by Frank et al. (2015), Nelson, Karoui, et al. (2017) and our study, most notably the type and modality of stimuli used (visually presented sentences vs. auditory narratives) and recording modality (scalp recorded MEG and EEG vs. intracranial recordings). A possible explanation for the divergent findings with respect to the study by Frank et al. (2015) is that measuring neural responses in settings which more closely resemble those of the brain’s natural mode of operation, brain dynamics will more readily reflect context-driven computations (captured by word entropy) which would explain why we find an effect of entropy in our dataset, but Frank et al. (2015) do not.

Second, different from Nelson, Karoui, et al. (2017), we do not observe significant modulations of high gamma power in response to changes in word entropy. A plausible reason is that high gamma responses can be more readily estimated in responses to visual stimulation as opposed to auditory stimulation where signal modulation is dominated by speech-induced low frequency power. Further, compared to ECoG signals, the signal-to-noise ratio of MEG, particularly in the frequency range reported by Nelson, Karoui, et al. (2017), as well as the more limited spatial resolution, may have prohibited accurate estimation of high frequency components in the current experiment.

### 4.4 Limitations and future work

Three limitations to our presently adopted approach deserve to be mentioned. First, we used lexical probabilistic language models defined over individual words whereas our brain responses were recorded during comprehension of connected discourse (narratives). One may wonder whether *n*-gram models of linguistic sequences are at all adequate for investigating naturalistic brain responses. Specifically, *n*-gram Markov models reduce context-dependent linguistic computation by requiring only the past *n*−1 words for generating expectation over upcoming words whereas it has been shown in the past that humans exploit much wider contextual cues rapidly in online comprehension (e.g. Nieuwland & Van Berkum, 2006).

It is beyond the scope of the present work to resolve the apparent discrepancy between the simplicity of statistical models embodying the Markov assumption and their success in engineering and psycholinguistic applications (see Jurafsky & Martin, 2009, for an overview). We are currently planning further experiments to test and compare models of linguistic sequences that capture richer contextual and temporal dimensions.

Second, language models used presently were computed over actually observed word strings as opposed to parts-of-speech tags or other types of linguistic information. As such, we cannot restrict our claims to lexical processing per se as our perplexity and entropy values are in principle driven by various dynamics in linguistic structure (e.g. a word can be surprising because it occurs in an uncommon syntactic construction or because it occurs with words which are not semantically congruent). Our current approach is not able to differentiate between the effects of these different representational levels. Others have shown that there are detectable spatial and timing effects of syntactic information in addition to lexical information in fMRI (Brennan, Stabler, Van Wagenen, Luh, & Hale, 2016; Henderson et al., 2016) and time-domain EEG (Hale, Dyer, Kuncoro, & Brennan, 2018). When designing the study, we chose the simplest (sequential) model architecture known to be empirically successful in the past, and of which the representation is insensitive to biases from downstream analysis errors (e.g. by part-of-speech taggers or parsers) and biases from representational choices of linguistic abstraction layers. Future work by our group will focus on uncoupling the effects of syntactic and lexical probabilistic computation in source-level frequency-domain MEG signals.

Third, in our investigation of the spectral characteristics of brain dynamics we relied on a relatively coarse binary complexity contrast (i.e. high vs. low complexity bins) which was achieved by averaging entropy and perplexity per epoch, thus capturing “slow” fluctuations of the linguistic features aggregated across several words. The general model-based approach allows in principle for testing of parametric modulations in complexity and its relationship to brain responses using regression-based designs (e.g. as in Frank et al., 2015; Frank & Willems, 2017; Hale et al., 2018). Indeed, we initially attempted to build a GLM-type of analysis pipeline to leverage the variance in word-by-word linguistic features. Yet, our models estimated turned out to be difficult to interpret; we were uncertain whether these inconsistent results were due to implementation, analysis parameter choices, data quality, or a combination thereof. Given the exploratory character of the present study, we then conceived the current analysis pipeline sacrificing the granularity of language complexity metrics for the interpretability of the outcomes.

We believe the current study provides ground for further research by evaluating models against brain oscillatory activity along other possible dimensions, for example type of linguistic information exploited It by the model (e.g. lexical vs. syntactic), or by using different model architectures for processing sequence (e.g. recurrent neural networks). Especially interesting would be applications of recent multivariate techniques for extraction of the so-called predictive models or “temporal response functions” (Crosse, Di Liberto, Bednar, & Lalor, 2016) from naturalistic MEG data on the basis of model-based predictors that embody distinct underlying hypotheses.

### 4.5 General discussion

Uncertainty and probabilistic computation have received increased attention as viable paradigms for describing information processing at the cognitive (Chater, Tenenbaum, & Yuille, 2006) and neural levels (Hasson, 2017; Knill & Pouget, 2004). Whereas in domains such as decision-making there have been several proposals and experimental evidence on how probabilistic functions or variables can be implemented with neurophysiological variables (Pouget et al., 2013), we are currently lacking similar theoretical proposals in domains of language processing (see Armeni, Willems, & Frank, 2017, for discussion).

Probabilistic processing metrics represent a linking hypothesis between cognitive theories and observed brain signals (Brennan, 2016). Presently, the work employing these metrics (present study included) is not addressing the question of neural codes, that is, how probabilistic knowledge and functions needed for language understanding can be encoded and decoded with models of neural circuits. They serve, instead, in describing the basic phenomenon—a statistical relationship between cognitive variables and neurophysiological observables— which in itself requires explanation (see Armeni et al., 2017; Carlson, Goddard, Kaplan, Klein, & Ritchie, 2017; Shagrir & Bechtel, 2017, for similar remarks).

What are the missing pieces then? On the one hand, there are generative circuit models of frequency-specific oscillatory activity (Kopell, Ermentrout, Whittington, & Traub, 2000; Roopun, 2008). This work provides understanding of conditions under which oscillatory activity can occur in brain circuits. It also outlines what computational roles can in principle be served by specific frequency bands, for example, that the beta band is suited for synchronization in circuits with long conduction delays for non-local computations (Kopell et al., 2000). On the other hand, oscillatory dynamics have been linked to general computational architectures for probabilistic (Bayesian) inference in neural circuits. For example, frequency contents of neural circuit activities could be mapped to specific computations for inference, e.g. feed-forward and recurrent information transfer between brain areas (Bastos et al., 2012, see Lewis & Bastiaansen, 2015, for an application of this account to language processing). Future work on human-recorded brain data will require a gradual move in emphasis from delineating correlational empirical phenomena towards explanatory links between known circuit properties in (human) electrophysiology and biologically-plausible computational architectures for language processing.

## 5 Conclusion

In the current study, we show that formalized probabilistic computations relate to frequency-specific brain dynamics during auditory narrative comprehension. We show that while theta-band dynamics relate to word-by-word contextual predictability likely reflecting lexical computations, beta-band dynamics are related both to word entropy and word perplexity, possibly reflecting maintenance of ongoing cognitive context which is updated upon encountering less expected words. Future challenges lie in using models that can exploit longer temporal relationships between words in narratives and in estimating predictive computations over different levels of linguistic representation.

## Acknowledgments

This work was supported by the Netherlands Organization for Scientific Research (NWO) (VIDI Grant 276-89-007 to R. M. Willems and VIDI Grant 864-14-011 to J. M. Schoffelen). The authors would like to thank Ashley Lewis for valuable comments on a previous version of the manuscript.

## A

fn001078

Met welk voertuig komt de verteller aan?

a) Een boot
b) Een vliegtuig
c) Per trein

Answer: b)

Aan welke rivier ligt de stad waar hij aan komt?

a) De Amazone
b) De Mississippi
c) De Rijn

Answer: a)

fn001155

In welk landschap bevindt de hoofdpersoon zich?

a) De jungle
b) Rivierdelta
c) Woestijn

Answer: c)

Wat voor een natuurramp heeft plaatsgevonden?

a) Een orkaan
b) Een aardbeving
c) Een tsunami

Answer: b)

fn001293

Welke dieren ziet de hoofdpersoon aan de bosrand?

a) Herten
b) Elanden
c) Wolven

Answer: a)

Hoe is het weer in het verhaal?

a) Zeer warm
b) Koud
c) Wordt niet beschreven

Answer: b)

fn001443

Waar bevindt de hoofdpersoon zich?

a) In de bergen
b) In de woestijn
c) Op een eiland

Answer: c)

Hoe heet de afwezige vriend van de hoofdpersoon?

a) Harald
b) Jonathan
c) Simon

Answer: b)

fn001498

Welk dier brengt ongeluk volgens het verhaal?

a) Raaf
b) Vleermuis
c) Uil

Answer: c)

Waar is een van de personages mee gestopt?

a) Roken
b) Drinken
c) Gokken

Answer: a)

## SUPPLEMENTARY INFORMATION

### Epoch creation respecting word onsets

We repeated the analysis with a different way of epoch creation compared to the procedure described in section 2.5 of the manuscript. Instead of creating 0.5 second data snippets (with 0 % overlap), without considering the word boundaries, we created epochs which were explicitly locked to the onset of words. In this way, epoch onsets did not straddle word boundaries, but this strategy resulted in highly overlapping data snippets. To allow for a fair comparison to the original analysis results, we subsequently removed the epochs with a high amount of temporal overlap, allowing for a maximal 20% overlap between the epochs.

We show the outcome of this analysis in the figures below. This analysis largely reproduces the patterns obtained with the original pipeline. We notice that the overall spatio-temporal extent of clusters (i.e. observed data patterns after thresholding) is less pronounced. E.g. the originally reported medio-lateral positive theta cluster for entropy contrast (Figure 4, panel A, of the manuscript) is now clustered by the algorithm into two separate clusters, one medial and a separate lateral cluster (rows A and B, Figure S2, respectively); the entropy-beta band differences are clustered to a smaller extent (row C, Figure 2). Similar observations hold for the perplexity contrast (Figure 2).

The overall contrasts (perplexity, entropy) in this pipeline are not conservatively controlled for the family-wise error rate at the desired alpha-level of 0.05.

**Figure S1.**
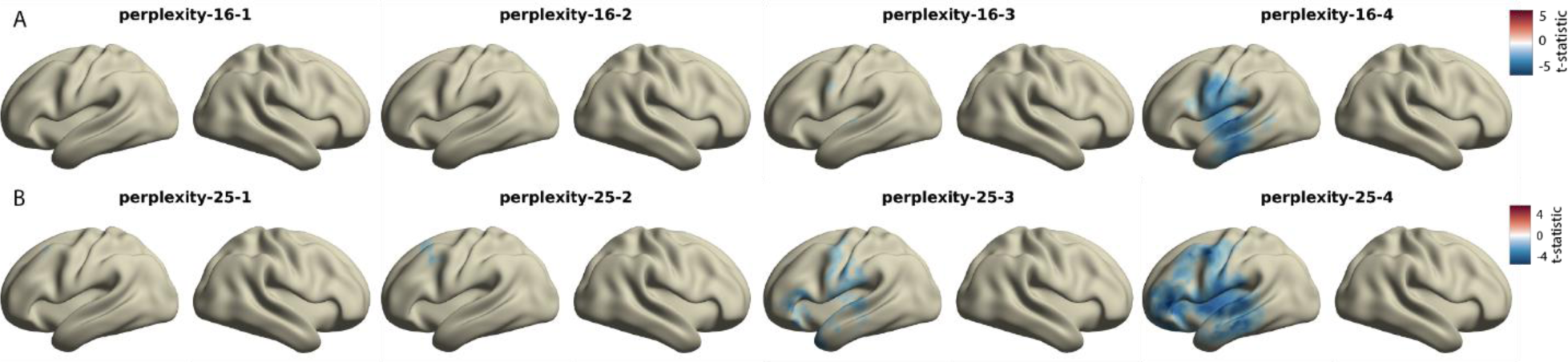
Group level t-maps for low beta (11--21 Hz, row A) and high beta band (21--31 Hz, row B) power quantifying differences between high and low perplexity bins. Source maps show paired-samples t-statistic per source location and time lag (0, 200, 400, and 600 msec). For the purposes of visualization, displayed are only t-statistics for source locations belonging to the cluster with the highest negative cluster statistic, that is, source locations not belonging to the maximal clusters the cluster are set to a value of zero.

**Figure S2.**
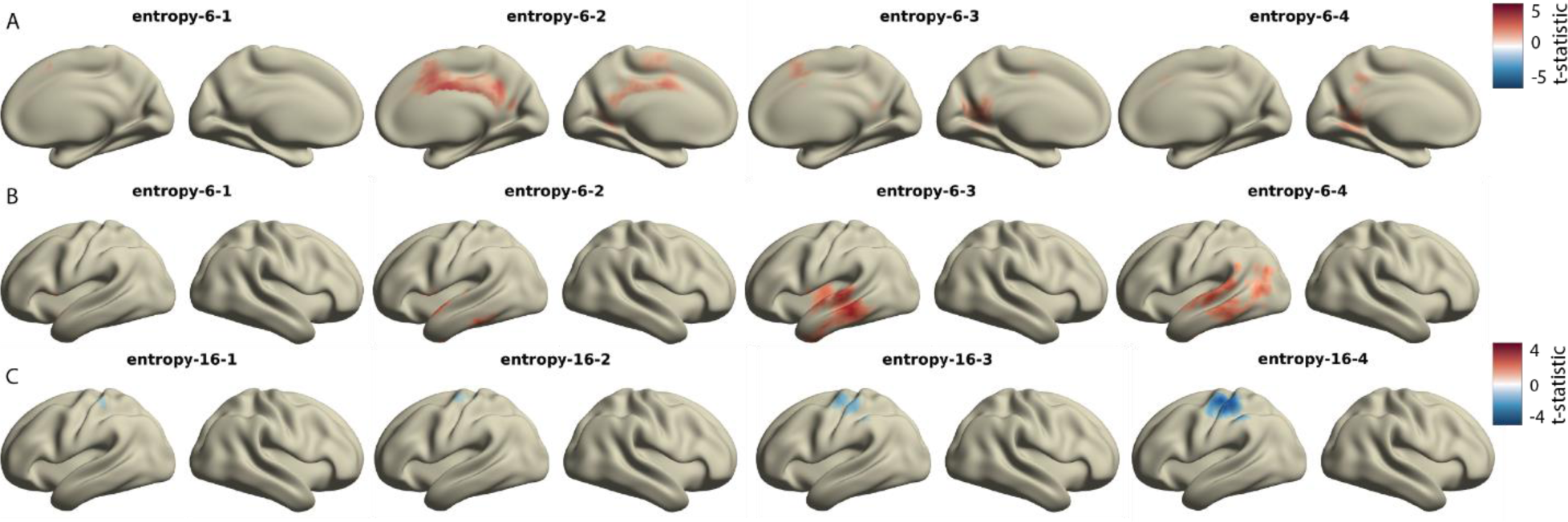
Frequency-specific power differences between conditions visualized here as group-level t-maps for theta-band (4--8 Hz, rows A and B) and lower beta-band (11--21 Hz, row C) power quantifying differences between high and low entropy bins. Source maps show paired-samples t-statistic per source location and time lag (0, 200, 400, and 600 msec). We show the first two highest positive clusters and the highest negative cluster for theta and beta bands, respectively. For the purposes of visualization, displayed are only t-statistics for source locations belonging to the cluster with the highest cluster statistic, that is, source locations not belonging to the maximal clusters the cluster are set to a value of zero.

